# Molecular subtyping and prognostic assessment based on tumor mutation burden in patients with lung adenocarcinomas

**DOI:** 10.1101/553461

**Authors:** Changzheng Wang, Fuqiang Li, Guoyun Xie, Sitan Qiao, Xulian Shi, Jianlian Deng, Han Liang, Cong Lin, Xin Zhao, Kui Wu, Xiuqing Zhang

## Abstract

Tumor mutation burden (TMB) is a potential biomarker for response to immunotherapy. The subset of patients with TMB has not been well characterized in lung adenocarcinomas. Here we performed molecular subtyping based on TMB and compared the features of different subtypes including clinical features, somatic driver genes and mutational signatures. We found that patients with lower tumor mutation burden had a longer disease-free survival, while higher tumor mutation burden is associated with smoking and aging. Analysis of somatic driver genes and mutational signatures demonstrates a significant association between somatic *RYR2* mutations and the subtype with higher mutation burden. Overall, our study identified two molecular subtypes based on TMB and described the corresponding difference in their clinical and genomic levels.

## Introduction

Lung cancer is the most frequently diagnosed cancer and one of the leading causes of cancerous deaths globally [1]. The estimated new cases of lung cancer were 222500 and the number of estimated death was 155870 in USA 2017 [2]. In China 2015, the age standardized incidence rate of lung cancer was 733300 and the estimated mortality was 610200 [3]. Adenocarcinoma is one of the most common histologic types of lung cancer. Cancer is a complex disease caused by the accumulation of genetic alteration and genome instability [4]. Many endogenous and exogenous factors, such as DNA damage repair inactivation, DNA replication mismatch, microsatellite instability and carcinogen exposure, can lead to increased somatic mutations. The total number of mutation occurred in DNA sequences of tumor cells is termed the tumor mutation burden (TMB) to sketched out the status of genomic mutation [5]. Notably, tobacco smoking is the major cause of lung adenocarcinoma with a high mutation burden [6].

TMB is an emerging biomarker for response to immunotherapy, since higher mutation burden is likely to harbor more neoantigens as targets of activated immune cells. The positive relationship between TMB and response to CTLA-4 and PD-1 inhibition has been show in melanoma and non-small cell lung cancer [7,8]. The increased frequencies of base-pair mutations have been described as one form of genomic instability which is an evolving hallmark of cancer [9]. TMB, as an index of genomic instability, may also be a biomarker for characterizing patients who were not treated by immunotherapy. The Cancer Genome Atlas (TCGA) project have used whole exome sequencing (WES) to measure TMB across 30 cancer types [10]. However, a precise understanding of the TMB in lung adenocarcinoma is still limited. Therefore, we hypothesize that mutation burden indicates the degree of genomic instability and may be correlated with clinical features in lung adenocarcinoma.

To this end, we performed a molecular subtyping based on TMB and analysis the difference between high tumor mutation burden (TMB-H) and low tumor mutation burden (TMB-L) from the aspects of clinical features, somatic driver genes and mutational signatures. Our analysis found that TMB may be a potential prognostic assessment marker and identified the significant difference between *RYR2* mutations of TMB-H and TMB-L. Furthermore, unbiased enrichment analysis also showed that *RYR2* associated with signature 4, which was predominant in TMB-H.

## Materials and Methods

### lung adenocarcinoma genome data

All the somatic mutations data, transcriptome sequencing data, and clinical information were downloaded and collected from previous published Chinese lung adenocarcinoma project [11]. This project hereinafter will be referred to LUAD_BGI.

### Mutation burden cluster

Considering the instability of outlier in the LUAD_BGI somatic mutation data, we used dynamic programming to cluster univariant data given by mutation burden of all the samples into optimal groups. This algorithm guarantees the optimality of clustering based on the minimum within-cluster sums of squares to give the optimal number of clusters k. To test the cluster number k, a range from 1 to 9 is provided for k and the optimal number of clusters was determained by Bayesian information criterion. The R package, Ckmeans.1d.dp [12], was perfromed for selecting the optimal number of clusters k.

### Identification of somatic driver genes

We predicted somatic driver genes using iCAGES [13], an efficient tool to search cancer driver genes, based on somatic mutation data of each case. Three layers of analysis steps were executed in iCAGES tool. In the first layer, Support Vector Machine was trained on somatic single nucleotide variants from COSMIC and Uniport databases to calculate SVM score and evaluate the driver mutation potential of each mutaiton. The second layer weights each mutation through integrating candidate mutation from first layer with prior biological knowledge on genetic-phenotypic assocaiation information. The third layer prioritizes for candidate driver muatation with corresponding drug activity score from PubChem database. For the identified somatic driver genes, they were selected from iCAGES candidate gene list if the gene was regarded as a driver gene in at least 5 cases.

### Mutation signature analysis

The mutation signature analysis is a procedure of deconvoluting cancer somatic mutations counts, stratified by mutation contexts or biologically meaningful subgroups, into a set of characteristic signatures and inferring the activity of each of the discovered signatures across samples. All SNVs were classified into 96 possible mutation categories based on the six base substitutions (C>A, C>G, C>T, T>A, T>C, and T>G) according to complementary base-pairing and 16 possible combinations of neighboring bases within the trinucleotide sequence context. We used SignatureAnalyzer, which uses a Bayesian variant of NMF and was recently applied to several cancer genome projects [14,15].

### Permutation test of signature enrichment analysis

The correlation between the activity of signatures and the overall tumor mutation burden could confuse the relationship of genes and the associated signatures. A traditional statistical test which compares the activity of signature among samples where the gene is wild type versus mutant for searching discrepant genes overestimates the significance of P values relatied to tumor mutation burden. The samples with a higher mutation burden prefer to generate more mutations which confound the statistical test power.

We controlled both the gene-specific and sample-specific mutation counts during the random permutation process to generate gene x sample binary mutation matirx, following the ‘Curveball algorithm’ described by Strona et al. [16]. We used one-tailed Wilcoxon rank-sum test to compare the signature activity of wild and mutant samples of a given gene. The statistic T_observed_ was a Wilcoxon statistic of actually observed data; the statistic T^r^_random_ was a Wilcoxon statistic of every permuted binary mutation matrix, where r=1, 2,…, 200000, the total number of permutation times. The finial P value of a given gene was the fraction of T^r^_random_ more extreme than T_observed_. This permutation test was followed by the discription from Kim et al. [14].

### Gene expression analysis

The gene expression level was calculated by using RPKM method[17] from 56 lung adenocarcinomas in LUAD_BGI. Differential gene expression was determined by edgeR, a Bioconductor package[18], between high mutation burden and low mutation burden groups

## Result

### Molecular subtyping based on mutation burden

Across the data from LUAD_BGI, the values of tumor mutation burden vary from 7 to 1126 with a mean value 163.5 (Fig. 1A). To determine the critical value of TMB, we took the routine method into account, that population is always divided into groups by mean or median values. For further consideration, we used a dynamic programing optimal univariate k-means clustering method to choose a better grouping threshold between mean and median values. The groups divided by the mean of TMB resembles the cluster result of optimal univariate k-means clustering (Fig. 1B). Finally, 33 patients were classified into TMB-H group, while the other 68 patients classified into TMB-L in accordance with the mean of tumor mutation burden.

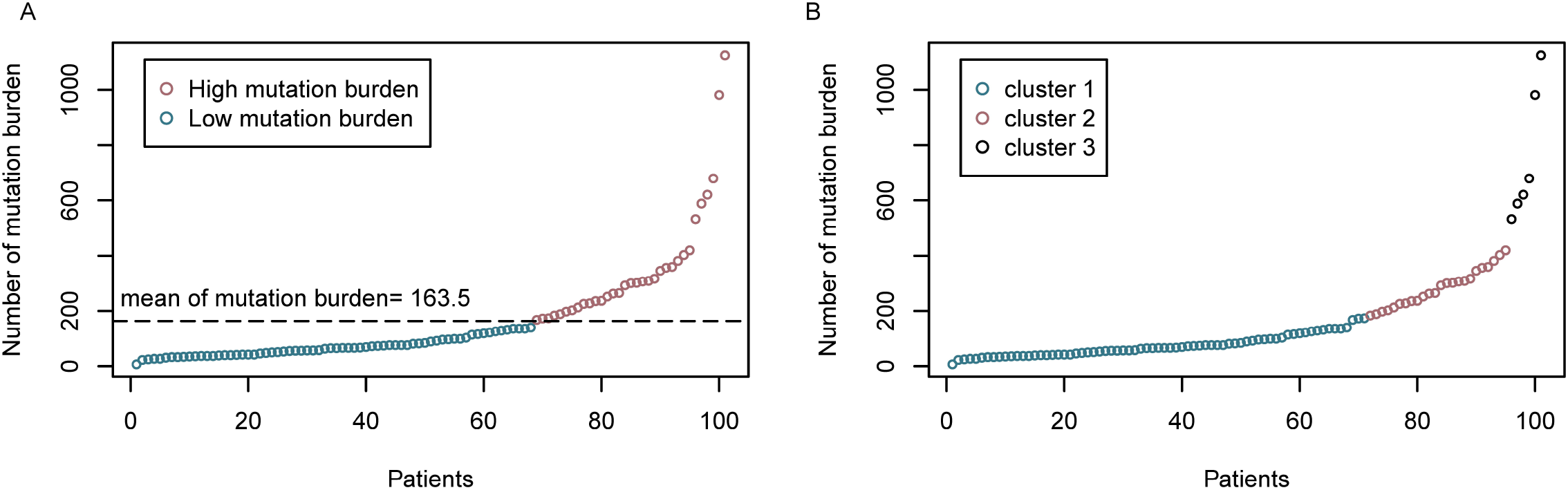

### Somatic driver genes from personal mutation background

Somatic mutations, including nonsense, missense, splice site, synonymous, frame-shift indel and nonframe-shift indel, were dectected in 33 cases and 68 cases in TMB-H and TMB-L respectively. Tumor mutation burden is derived from the genes harbored mutations of each patient. To determine the individual mutation profile, we used iCAGES to identify somatic driver genes based on somatic mutation background and mutation burden status of each case as described in the method section. The comprehensive scores of candidate somatic driver genes were evaluated based on their mutational, fuctional, and drug actionable characteristics.

In the total 101 cases of LUAD_BGI cohort, 26 somatic driver genes were identified and mutated in at least 5 cases. In those somatic driver genes, 9 genes, *TP53*(40%), *EGFR*(28%), *CDC27*(20%), *KRAS*(14%), *LAMA2*(8%), *PIK3CA*(7%), *TRIO*(7%), ATR(6%) and *BRAF*(5%) were previously reported, whereas the other 17 genes, *RYR2*(29%), *COL11A1*(13%), *HERC2*(11%), *LRP2*(11%), *SI*(11%), *RELN*(9%), *ITGA8*(9%), *UBR4*(8%), *HTT*(7%), *ADCY2*(7%), *COL5A1*(7%), *FGA*(7%), *GRM1*(7%), *GLI3*(6%), *TSHR*(6%), *GRIA1*(5%) and *SCN5A*(5%) were not reported previously (Fig. 2). In particular, *TP53, EGFR, RYR2, KRAS* and *CDC27* were identified as driver genes in more than 10 cases using ICAGEs. Importantly, only *RYR2* was reported for the first time as the mutated cancer driver according to Cancer Gene Census [19] and IntOGen [20].

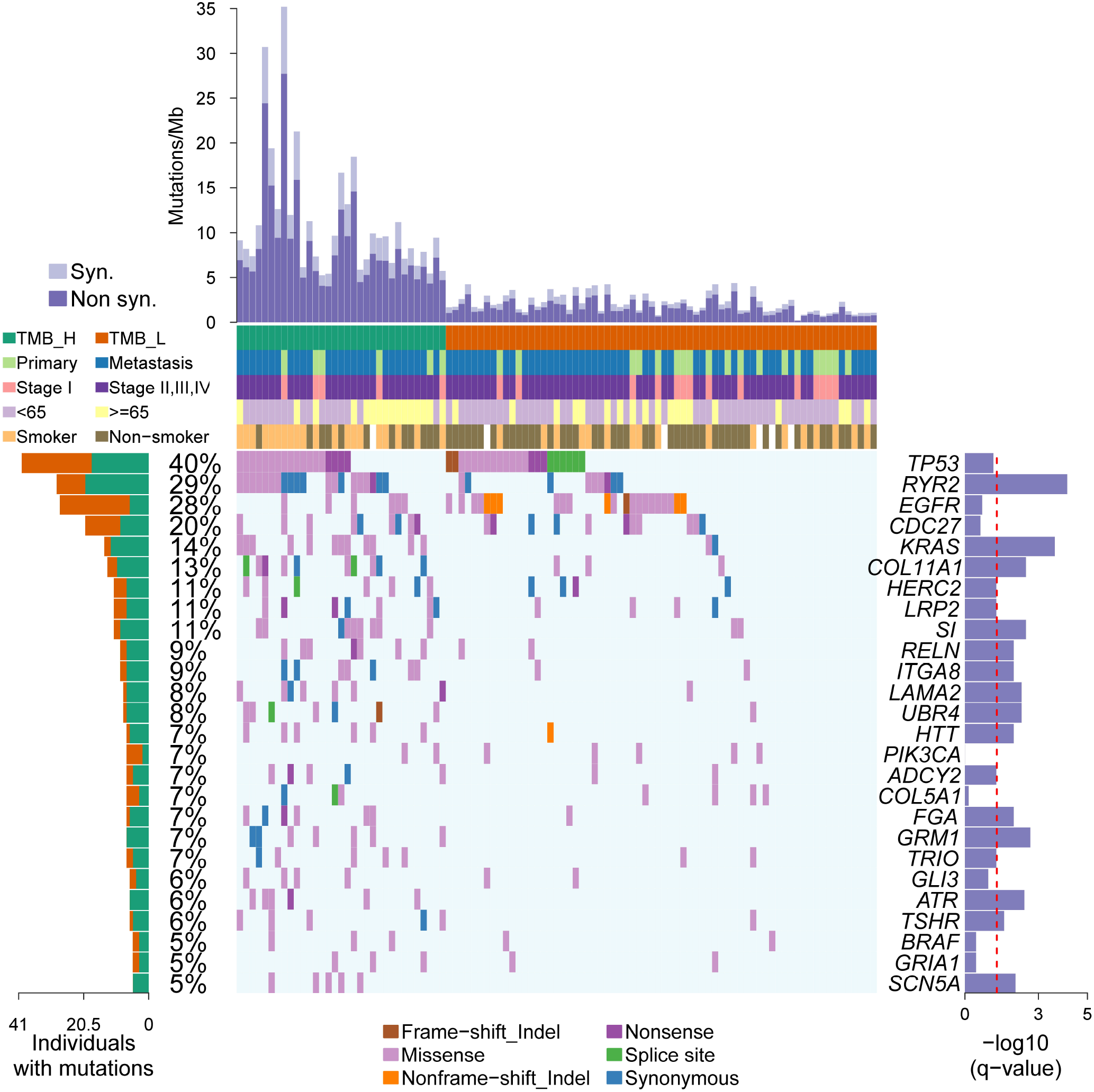

To determine the driver genes differential between TMB-H and TMB-L, two identified molecular subtyping groups, fisher exact test was performed to figure out the significant driver genes. BetweenTMB-H and TMB-L groups, 14 genes were showed significant difference and enriched in TMB-H groups. The mutational tendency of *EGFR, CDC27* and *PIK3CA* revealed a potential feature in TMB-L groups, although these genes were not significant different between TMB-H and TMB-L groups.of Among these 14 genes, *RYR2* was the most significant gene with a q value of 6.69e-5 indicating the overrepresentation of mutated *RYR2* in TMB-H groups.

### Clinical features associated with tumor mutation burden

Next, we investigated the influence of clincal freatures related to tumor mutation burden. We integrated clinical features of the 101 cases into aforementioned molecular subtypes to explore whether the clinical features were associated with the two TMB groups. Among the clinical factors of age, metastasis status, smoking historyand tumor stage, only smoking history and age were assocaited with TMB levels. There were significantly more patients over age 65or with smokig history in TMB-H group with the p-value of 0.0006 and 0.042 respectively. Higher percentages of patients who were under worse status, such as late tumor stage and tumor metastasis condition, were also observed in TMB-H group, indicating these clinical factors may associate with higher TMB levels in some degrees(Fig. 3A). Further analysis of the risk factors, aging and smoking, showed a greater number of TMBs in patients older than 65 (p-value=0.0141) or with smoking history (p-value=0.0009) (Fig. 3B, 3C). The disparity of the number of TMB in comparison of age>=65 and age<65 groups, and smoker and non-smoker groups, suggest that the accumulation of TMB in age>=65and smoker group is much higher than age<65 group and non-smoker group. Further, Kaplan-Meier survival analysis showed a significantly longer disease-free survival (DFS) in the patients with low TMB levels after surgery (p-value = 0.0133, Fig. 4). This result suggests that TMB is potentially associated with surgical outcome in lung adenocarcinoma.

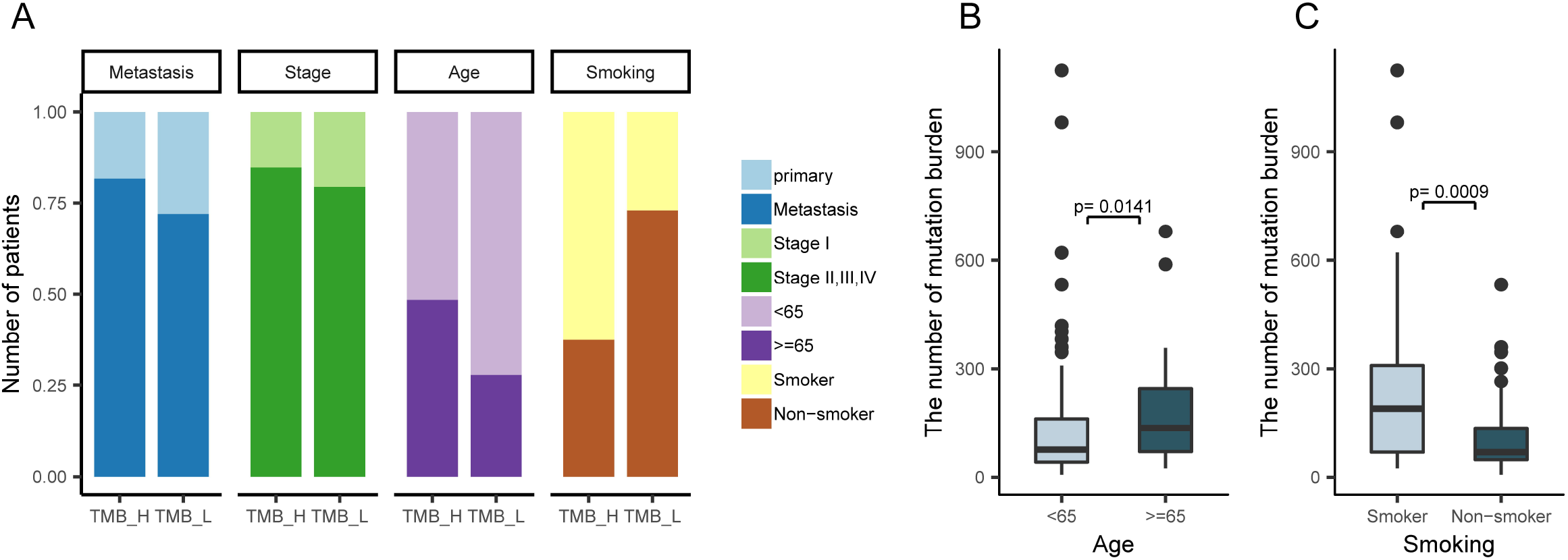

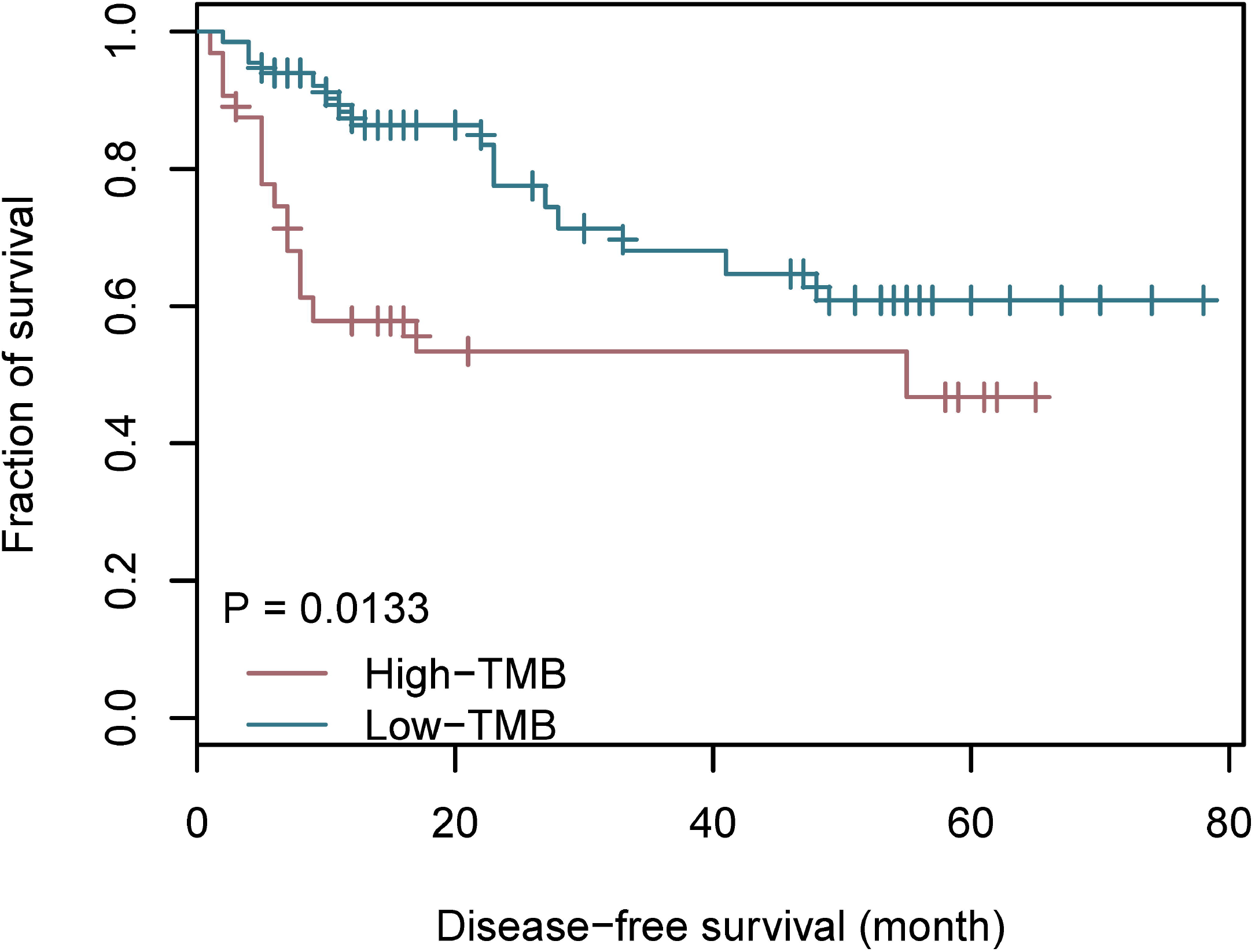

### Mutation signature analysis of lung adenocarcinoma in tumor mutation burden subtypes

To understand the mutation accumulation processes during lung adenocarcinoma, we performed mutational signature analysis on the 101 cases. We applied a Bayesian NMF algorithm to mutation counts, stratified by 96 trinucleotide mutational contexts, to infer (i) the number of operating mutational processes, (ii) their signatures (96 normalized weights per process), and (iii) the activity of each signature in lung adenocarcinoma (the estimated number of mutations associated with each signature) [14].

Our analysis identified three mutational signatures in LUAD-BGI cohort, and we compared them with those applied by the Sanger Institute, which are described in the COSMIC database (http://cancer.sanger.ac.uk). Signature W1, characterized by C>G transversions and C>T transitions at TpCp[A/C/G/T] motifs, is corresponding to COSMIC signature 2 (cosine similarity of 0.85), which also attributed to activity of the AID/APOBEC family of cytidine deaminases (Fig. 5A). Signature W2, charecterized by C>A transversions at a broad spectrum of bases context, was closely resembles COSMIC signature 4 (cosine similarity of 0.96). COSMIC signature 4 is associated with smoking and its profile is similar to the mutational pattern observed in experimental systems exposed to tobacco carcinogens (e.g., benzo[a]pyrene). Signature W3, characterized by C>T transitions at [A/C/G]pCpG motifs, is corresponding to COSMIC signature 6 (cosine similarity of 0.91) that is associated with defective DNA mismatch repair.

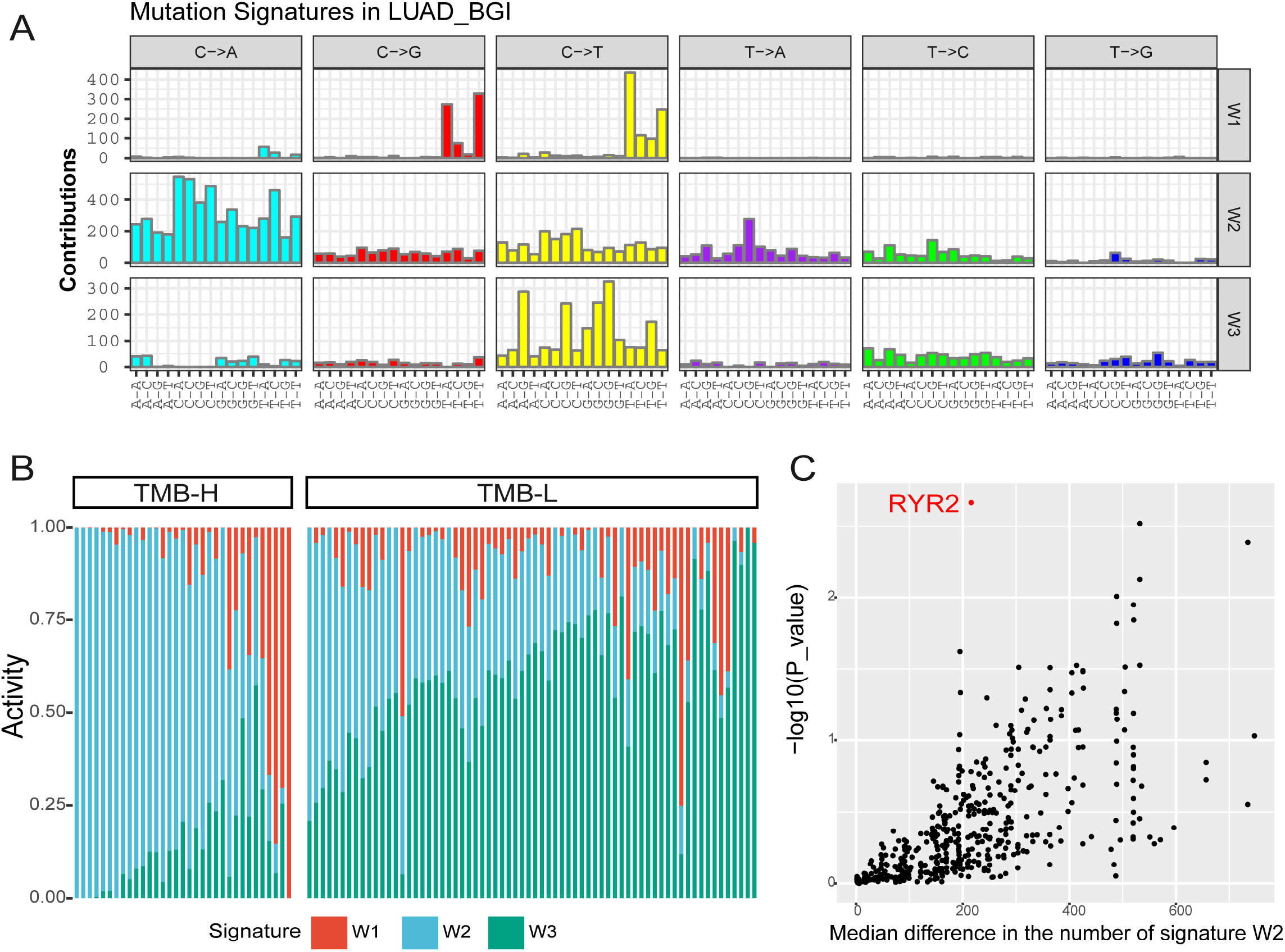

The TMB-H and TMB-L groups were clearly distinguished by mutation signature spectrums, espcially the activity of signature 4 (Fig. 5B), suggesting a underlying impact of mutation signature on molecular subtyping.

### Signature 4 activity is associated with *RYR2* mutations

We performed signature ehrichment analysis to further characterize mutated genes that associated with signature 4. We compared the activity of signature 4 in tumors that harbored a nonsynonmous mutation in the gene and tumors that did not cross the LUAD_BGI cohort. To exclude the noise resulted from tumor mutation burden, we assessed the significance level using a permutation-based method that contrals the tumor mutation burden in each sample [14]. We found that *RYR2* was the top significant gene associated with siganature 4 acitivity (Fig. 5C).

### Profile of gene expression level in tumor mutation burden subtypes

Unsupervised hierarchical clustering analysis was performed based on 212 differential genes in 56 samples (supplementary data), 19 of which was from TMB-H and 37 samples from TMB-L. We identified two mRNA groups with distinct mutation burden characteristics, which correspoded to mutation burden subtypes with only one sample of TMB-H clustered to TMB-L (Fig. 6). We also found that a gene cluster with 19 genes (*FMN1, FHIT, FDXR, HHLA2, F2RL1, PLXNA2, TNKS1BP1, CACNB1, SPINK5, BTBD9, CRYM, MPV17L, SH3RF2, SULT1C2, ABCC3, FCGBP, ST6GALNAC1, CLDN1, GDF15*) were down regulated in TMB-H. Using gene set enrichment analysis, we identified that these genes were significantly enriched (FDR<0.05) with a gene set which was down-regulated in epidermis after to UVB irradiation associated with mutations accumulation[21].

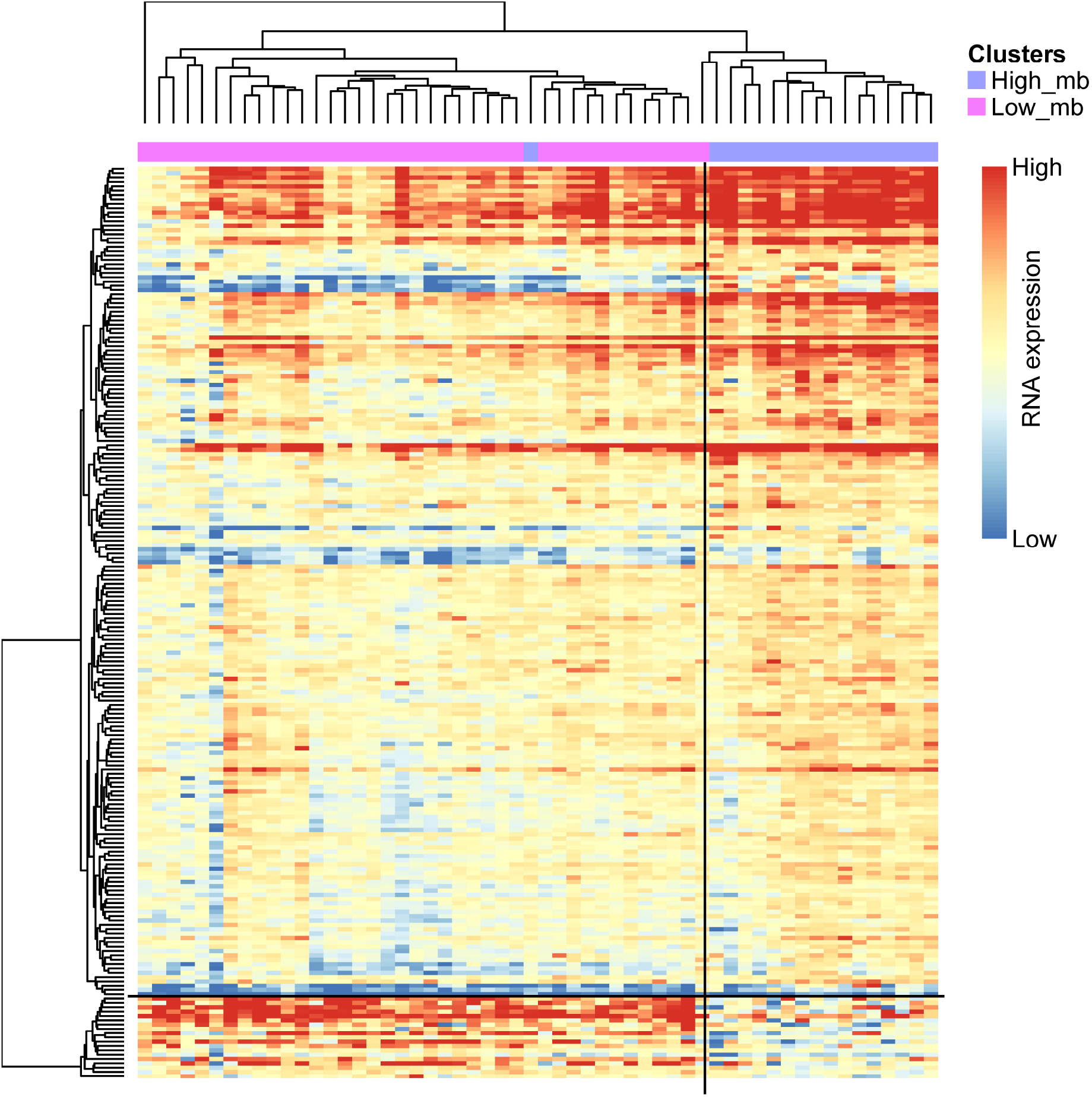

## Discussion

We constructed a molecular subtype model based on the tumor mutation burden and showed that the patients from LUAD_BGI cohort can be further stratified into TMB-H and TMB-L subtypes by the mean of population TMBs, which was analogous to the result of dynamic programing optimal univariate k-means clustering method. Patients in the subset of TMB-L had a better prognosis as evidenced by a longer DFS compared with patients in TMB-H subset. Tumor mutation burden as a genomic marker of prognosis and predictor of treatment response has been reported in ovarian cancer [22]. In this study, for the first time, we demonstrated that lower tumor mutation burden predicts more favorable DFS in patients with lung adenocarcinoma. The molecular subsets of TMB-L and TMB-H could assist the assessment of prognosis for further clinical research of lung cancer. Exploring the clinical features associated with tumor mutation burden is also essential to understand the molecular subtypes. The higher TMB was significantly associated with the group of patients elder than 65 or with smoking history, which were the risk factors of carcinogenesis in a common sense [23]. TMB-H group consisted of patients associated with cancer risk factors is in accord with the result of unfavorable prognosis. There were mutually verifying relationships between the results of survival analysis and clinical features association analysis. Compared with other genome-based molecular subtyping studies [24–26], this finding highlighted the importance of molecular subtyping based on mutation burden in clinical utility.

However, a recent research demonstrated that high TMB was associated with a better prognosis in patients with resected non-small-cell lung cancer (NSCLC), while lung cancer-specific survival with adjuvant chemotherapy was more significant in patients with low TMB[27]. We speculate that one of the possible explanations could be the different regions of TMB result in the different association between TMB and prognosis, and in our study, TMB was calculated by nonsynonymous mutations in whole exon regions, not targeted panels. These observations and underlying mechanisms between TMB and prognosis should be confirmed by further studies.

In addition to the association with clinical features, we also found that signature 4 activity was increased in TMB-H group using signature ehrichment analysis. Signature 4 is associated with smoking which reflects that one of the clinical features TMB-H also associated with was smoking. This signature was found only in cancer types in which tobacco smoking increases risk and mainly in those derived from epithelia directly exposed to tobacco smoke [26]. The inter relations with smoking demostrated that the mutation burden triggered by tobacco exposure might be detected as the profile of signautre 4 in TMB-H. The molcecular subtypes of lung adenocarcinomas can be described as the pattern of the TMB-H enriched with signature 4 and TMB-L. Besides, we identified an association between Signature 4 activity and somatic mutations in *RYR2*, which was also mutated signigicantly in TMB-H, providing the important insights for molecular subtyping based on tumor mutation burden.

*RYR2*, Ryanodine Receptor 2, encodes a ryanodine receptor found in cardiac muscle sarcoplasmic reticulum and induces the release of calcium from the sarcoplasmic reticulum into the cytosol [28]. Mutations in *RYR2* are frequently reported in ventricular tachycardia [29–31] and Arrhythmogenic right ventricular dysplasia type 2 [32]. Although *RYR2* is known to associate with heart disease, *RYR2* as one of the components of a calcium channel also influences calcium signaling in airway smooth muscle cells [33–35]. Recent research demonstrated the association of *RYR2* with asthma by genome-wide analysis [36]. Cigarette Smoke exposure causes down-regulation of ryanodine receptor in mice airway smooth muscle and small airway contraction, which is a major site of airflow limitation in chronic obstructive pulmonary disease (COPD) [37]. Mutations in *RYR2* may result in alteration of airway smooth muscle and asthma which is a possible risk factor for lung cancer [38,39]. This is a possible underlying mechanism by which the association between *RYR2* and lung cancer could arise.

Together, our analysis suggests that molecular subtyping based on TMB could be treated as a prognostic marker. Also, signautre 4 and the significance of mutation in *RYR2* are highlighted in TMB-H group. Further studies will be needed to characterize the mechanism underlying mutation of *RYR2* in lung cancer and to expolre potential relationships between *RYR2* and high tumor mutation burden.

## Acknowledgement

This work was supported by the National Key Research and Development Program of China (No. 2016YFC0902301). The funders had no role in study design, collection, analysis and interpretation of data, and in the writing the manuscript.

