## supplementary data for "Molecular subtyping and prognostic assessment based on tumor mutation burden in patients with lung adenocarcinomas"

Difference expression genes between high mutation burden and low mutation burden groups

*ABCC3, AFAP1L1, ANAPC5, ANKRD52, ATAD2, ATP5A1, ATP5B, AURKA, AURKB, BCCIP, BMP6, BNIP3, BRAP, BRIX1, BTBD9, BUB1, BUB3, BZW1, C10orf46, C12orf29, C12orf73, C15orf23, C18orf55, C6orf115, C9orf30, CACNB1, CACYBP, CBS, CBX3, CCDC102B, CCDC99, CCNA2, CCNB1, CCNB2, CCT4, CCT5, CCT7, CDC123, CDCA3, CDK1, CDK6, CDKN3, CENPO, CHAF1A, CKS1B, CLDN1, CNBP, CPSF3, CRYM, CUL2, CXXC1, DNAJC7, DNAJC9, DPM1, DSN1, DUSP4, DYNLL1, E2F1, ECT2, ERLIN1, ERMP1, EZH2, F2RL1, FAM175B, FAM204A, FANCG, FANCI, FBXO18, FCGBP, FDXR, FGFR1OP2, FHIT, FMN1, FNTA, FOXM1, GABPB1, GDF15, GGH, GIMAP4, GIMAP7, GMNN, GNLY, GPN1, GSTCD, GVINP1, GZMA, GZMB, H2AFZ, HAUS1, HAUS6, HHLA2, HLTF, HMGB2, HNRPLL, ING1, KCTD3, KIF11, KIF20A, KIFC1, KLRC3, LSM1, MAPKAP1, MAPRE1, MBD2, MCM2, MCM6, MED27, MEMO1, METTL4, MKI67, MMADHC, MPV17L, MRPL13, MRPL35, MRPL9, MRPS18C, MRPS22, MSH6, MTERF, MTHFD2, MYBL2, NCAPD2, NCAPG2, NDUFA12, NDUFA8, NDUFV2, NFYB, NKG7, NOL7, NRBP1, NUP37, NUSAP1, PAIP1, PDSS1, PHF19, PLEKHA8P1, PLK1, PLXNA2, POP7, PPP1R12A, PPP3R1, PRC1, PRIM1, PSMA4, PSMB7, PSMD14, PSRC1, PWP1, R3HDM1, RACGAP1, RANBP1, RBM17, REXO4, RFC4, RNF219, RPF1, RTKN2, SDAD1, SENP1, SEPHS1, SET, SH3RF2, SKP2, SMAD2, SMC2, SMC3, SMC6, SNRPA1, SNRPF, SNRPG, SPAG5, SPINK5, ST6GALNAC1, STARD7, STMN1, STOML2, SULT1C2, SUMO1, Symbol, TARS, TCF19, TEX10, TIAL1, TIMELESS, TIMM17A, TMEM194B, TMEM206, TNKS1BP1, TOMM5, TOPBP1, TP53INP1, TPRKB, TPX2, TTL, TXNL1, TXNL4A, UBE2S, UCK2, UFD1L, UHRF2, UNG, USMG5, VPS33B, VRK1, WDR5, WDR67, XPOT, YWHAQ, ZCCHC7, ZNF544, ZWILCH, ZWINT*
